# Vascular microforamina and endocranial surface: normal variation and distribution in adult humans

**DOI:** 10.1101/2023.12.15.571867

**Authors:** Emiliano Bruner, Stanislava Eisová

**Author notes:** Corresponding author: Emiliano Bruner, Centro Nacional de Investigación sobre la Evolución Humana, Paseo Sierra de Atapuerca 3, 09002 Burgos, Spain.

## Abstract

The term *craniovascular traits* refers to the imprints left by arteries and veins on the skull bones. These features can be used in anthropology and archaeology to investigate the morphology of the vascular network in extinct species and past populations. Generally, the term refers to macrovascular features of the endocranial cavity, like those associated with the middle meningeal artery, venous sinuses, emissary foramina, and diploic channels. However, small vascular passages (here called *microforamina* or *microchannels*) have been occasionally described on the endocranial surface. In this study, we describe and quantify the amount and distribution of these microforamina in adult humans (N = 45) and, preliminarily, in early to late juvenile subjects (N = 7). Adults display more microchannels than juvenile skulls. Females show more channels than males, and it should be evaluated whether this trait can represent a new described sexual features, influenced by sex biology. The distribution of the microforamina is particularly concentrated on the top of the vault, in particular along the sides of the sagittal, metopic, and coronal sutures, matching the course of major venous sinuses. Nonetheless, the density is lower in the region behind the *bregma*. A preferential pattern of distribution seems to join diagonally the coronal and dorsal parietal regions. Beyond oxygenation, these features are likely involved in endocranial thermal regulation and immune responses, and their distribution and prevalence can hence be of interest in human biology, evolutionary anthropology, and medicine.

## Introduction

The vascular system of the braincase is formed by a complex network of vessels running through the brain (cerebral vessels), the meninges (meningeal vessels), the cranial bones (diploic vessels), and the ectocranial surface (e.g., Scremin, 2004; Patel, 2009; Toriumi et al., 2011; Adeeb et al., 2012; Mortazavi et al., 2013; Tsutsumi et al., 2013). These vessels are generally labelled and assigned to a particular network according to their position and, in some cases, to some anatomical features. Nonetheless, they are part of a single widespread system, which allows the blood to move from one region to another in response to physiological changes and homeostatic conditions. Besides oxygenation, these vessels are also crucial for brain thermal regulation and hydrostatic support, two aspects that are essential for proper brain functioning. The term *craniovascular traits* generally refers to the imprints of these vessels within or onto the bone, and these features are employed in archaeology and anthropology to investigate vascular morphology in extinct species or past populations (Píšová et al., 2017; Rangél de Lázaro et al., 2018; Eisová et al., 2019). In fact, because of the importance of encephalization in hominid evolution, the distribution and prevalence of these features have been tentatively associated with evolutionary adaptations or physiological responses to brain heat production (Boyd, 1930; Kimbel, 1984; Falk, 1986, 1990, 1993; Bruner and Sherkat, 2008; Rangel de Lázaro et al., 2016). It has been also hypothesized that the complexity and topology of these vessels might be related to the balance between heat production and dissipation, which is influenced by changes in both brain size and shape (Bruner et al., 2011, 2012, 2014). In general, all these features deal with macroanatomical traits, which can be detected by visual inspection. However, the fractal nature of the vascular system generates an exponential number of small branches, which account for an important part of the blood flow and are more difficult to detect in osteological remains. For example, the analysis of the diploic channels can reveal and quantify the distribution of the large vessels running into the vault bones, but not of the small vessels running through the trabecular spaces (Rangel de Lázaro et al., 2020). It is hence likely that information on these minor channels is crucial to provide reliable estimates of the real distribution of the vascular networks. Actually, a large number of microvessels cross the endocranial surface, possibly bridging the diploic and meningeal networks, and influencing the thermal regulation of the endocranial space through the interaction with the cerebrospinal fluid (Zenker and Kubik, 1996). Recently, it has been also hypothesized that these small vessels might be involved in immune cell migration from the vault bones to the brain, in response to damage and inflammation processes (Herisson et al., 2018). Microforamina associated with these small vessels have been described in fossils dated to more than 800 thousand years (Bruner et al., 2017), as well as in Neandertals (Hui and Balzeau, 2023). However, the normal prevalence and distribution of these microforamina are not known for the modern population, as these traits are still scarcely investigated in the literature. In this survey, we analyse the presence and patterns of these microforamina in a sample of modern human skulls, to provide basic anatomical information for the study of these features.

## Materials and Methods

In this paper, we call *microforamina* or *microchannels* the scattered tiny vascular passages detectable on the endocranial surface, associated with small vessels (*microvessels*) bridging the diploic and the endocranial space.

The sample consists of Computed Tomography (CT) scans of crania of anatomically normal individuals selected from the Pachner collection, which is housed at the Department of the Anthropology and Human Genetics of Charles University in Prague. This is an autopsy collection from the 1930s formed by individuals belonging to the Czech urban working class. Contemporary documentation (age at death, sex) is available for most of the individuals of the sample (Borovanský, 1936; Pachner, 1937). Medical CT scans were obtained with a Siemens Somatom 16 (voxel size: 0.488 × 0.488 × 0.300 mm). The final sample contains 45 adult (17 males, 20 females, 8 unknown sex) and 7 juvenile skulls. Adult specimens range from 22 to 73 years of age (mean and standard deviation: 46 ± 14). The juvenile sample includes very different developmental ages and, also taking into account the small number of individuals, the corresponding results should be considered preliminary.

CT images were processed with 3D Slicer 4.10.2 and Mimics Innovation Suite 20.0. The detection of microforamina was performed in 2D views of the tomographic scans. The routes of the small channels running from the diploe to the endocranial surface can be visualized and followed by scrolling the stacks in the three planes, and the microforamina corresponds to the endocranial aperture of the channel. The diameter of the microforamina is usually smaller than 0.5 mm. In some cases the opening may be larger, up to 2 mm, approaching the size of the emissary veins. Three-dimensional coordinates of the microforamina were recorded, together with the coordinates of basic craniometric landmarks (*nasion, basion, lambda, asterion, porion, opisthion*). Standard skull diameters were also measured (*glabella*-*opisthocranion, eurion*-*eurion, basion*-*bregma*). To map the distribution of the microforamina in the whole sample, the cranial landmarks have been registered by Procrustes superimposition through *Morpheus et al*. (Slice, 2014). This registration translates, scales, and rotates the coordinate systems to minimise their spatial differences (Slice, 2007; Mitteroecker and Gunz, 2009; Zelditch et al., 2012). The number of the foramina in different groups has been compared through the Mann-Whitney test. Statistics were computed with PAST 4.12b (Hammer et al., 2001).

## Results

The number of microforamina does not show a normal distribution (Shapiro-Wilk p = 0.02), with most specimens displaying between 50 and 110 channels (Fig. 2). Table 1 shows the descriptive statistics (quartiles) for the number of microforamina in each group. Females show more channels than males, although this difference, in this sample, is not significant (p = 0.14). The difference between adults and juveniles is, instead, significant (p = 0.04). The number of microforamina is not correlated to age, cranial width, and cranial height. It is instead negatively correlated with cranial length (r = -0.31). In this sample, sexual cranial differences (larger size in males) are more evident for the cranial length (p < 0.0001) than for the width (p = 0.03) and height (p = 0.01). An analysis of covariance between the number of foramina (as covariate) and cranial length (as independent variable) adjusting for sex suggests that males and females show the same slope (p = 0.45) and the same adjusted mean (p = 0.90). Hence, the correlation between cranial length and the number of channels must be intended as associated with sexual differences.

**Table 1.** Number of microforamina.

|  | **N** | **25th** | **Median** | **75th** |
| --- | --- | --- | --- | --- |
| *Females* | 17 | 76 | 106 | 133 |
| *Males* | 20 | 69 | 88 | 113 |
| *All adults* | 45 | 73 | 93 | 118 |
| *Juveniles* | 7 | 40 | 72 | 86 |

**Figure 1.**
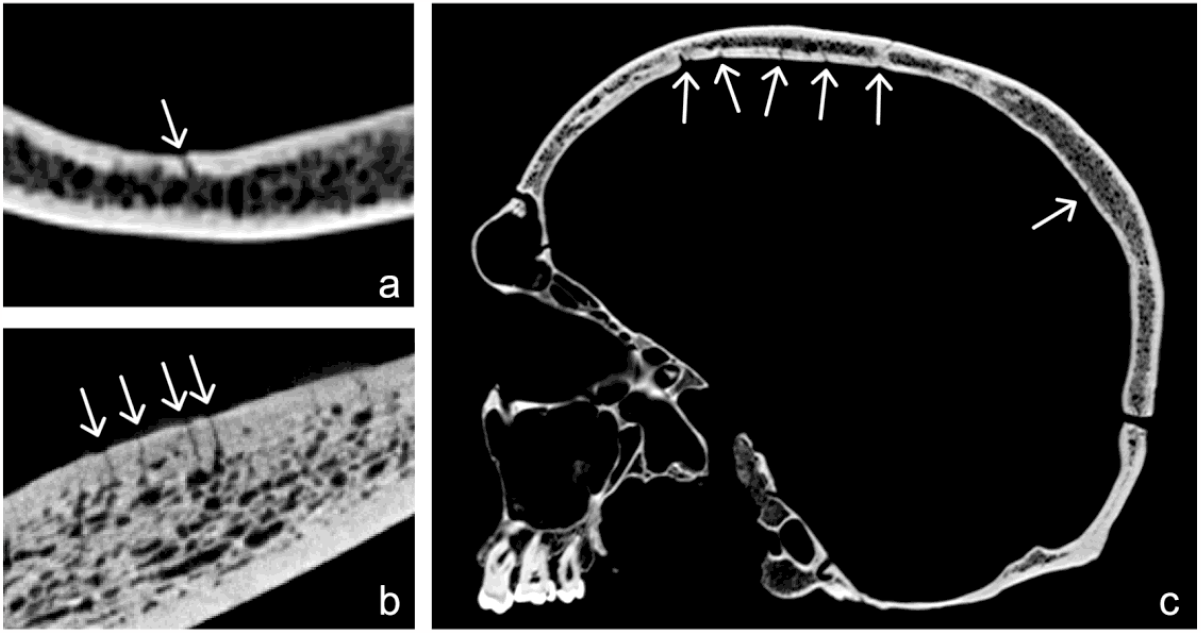
Microforamina on the endocranial surface (arrows), from a medical tomography (a), from a microtomographic scan (b), and in a skull from the current sample (c).

**Figure 2.**
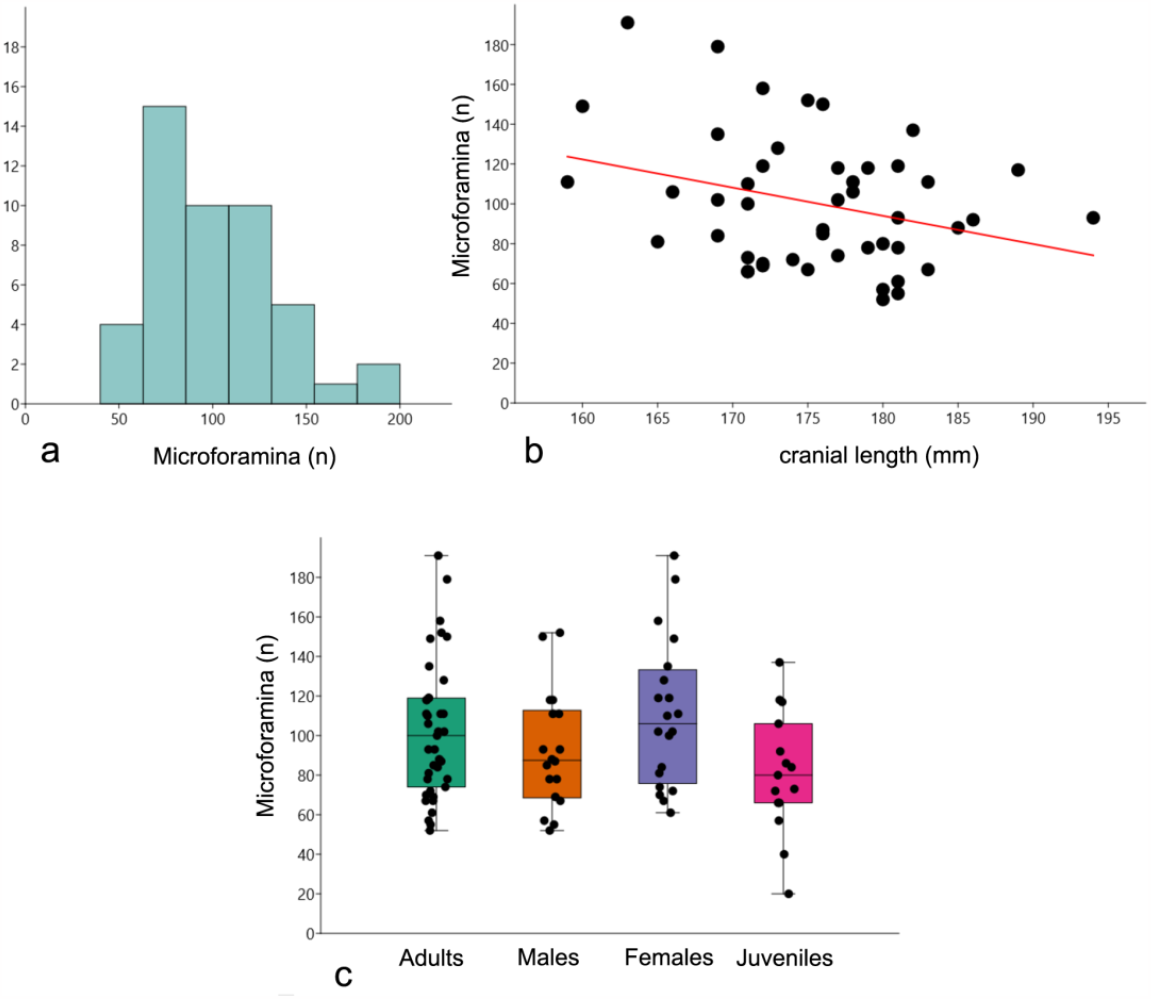
Distribution of the number of microforamina in the sample (a), correlation between cranial length (*basion*-*bregma*) and number of channels (b), and distribution of the number of channels in each group through jitterplot and non-parametric boxplots (c).

Usually, the microforamina are associated with small diploic veins, although they can be connected with the large diploic channels, the middle meningeal vessels, and the venous sinuses. They can be often detected in the area of the arachnoid foveae, remarking their implication in the flow of the cerebrospinal fluid. The endocranial distribution of the microforamina is shown, after the superimposition of the cranial landmarks, in Figures 3 and 4. Despite they are widespread throughout the whole neurocranial surface, the microforamina are particularly concentrated on the dorsal region of the braincase, and in particular along the sides of the sagittal, metopic, and coronal sutures. These regions match the course of the superior sagittal and sphenoparietal venous sinuses. Among these dorsal regions, the parietal squama looks the one with the lower density of channels, while the frontal bone displays a more homogeneous and dense distribution of microforamina. Interestingly, the region behind the *bregma* is an exception to this distribution, showing a lower channel density. Instead, a preferential line of distribution runs obliquely from the parasagittal region to the coronal region, before reaching this bregmatic area. Actually, the bregmatic region is apparently surrounded by a rhomboid pattern of channel density (Fig. 5), which suggests a spatial regularity due to underlying morphogenetic factors.

**Figure 3.**
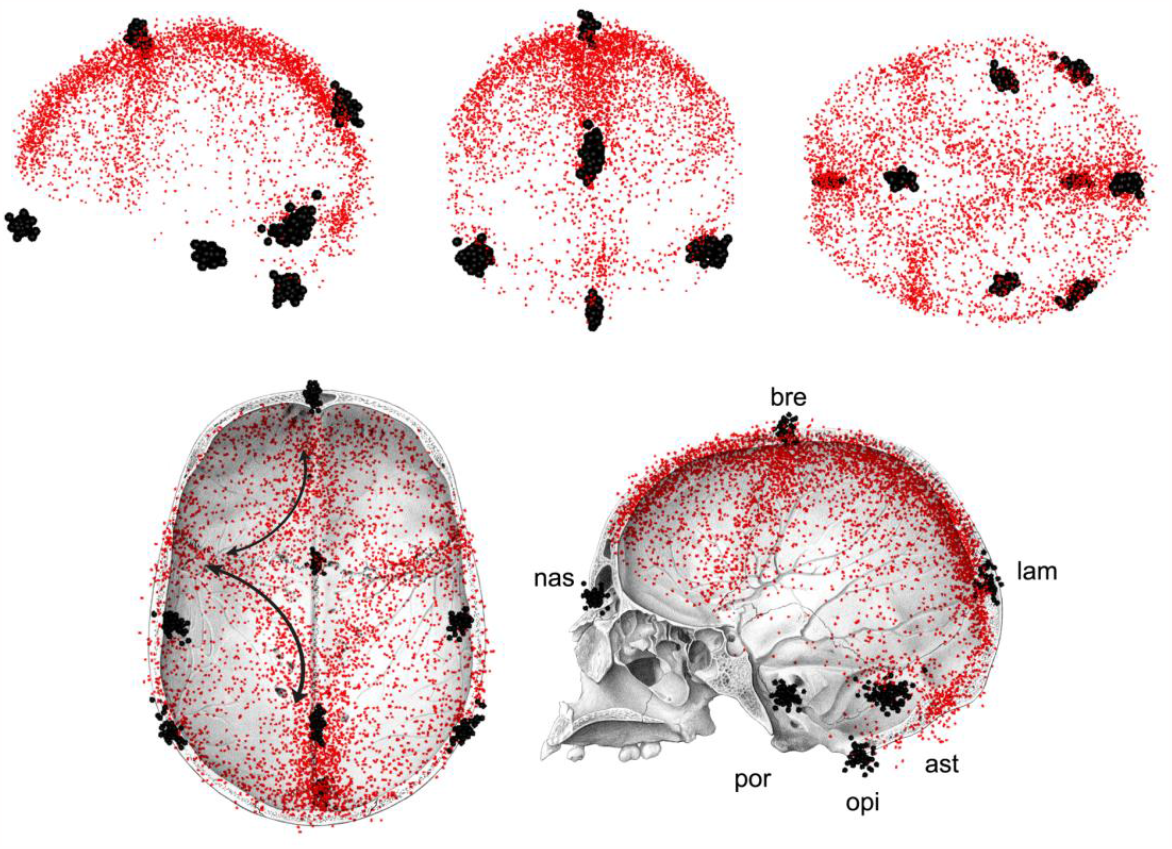
Above: maps of the distribution of the microforamina (red dots) after cranial superimposition (black dots) in the whole sample (left lateral, posterior, and superior view). Below: the same maps visualized on a drawing of a human skull (skullcap from inferior view and midsagittal lateral view). Labels: ast: *asterion*; bre: *bregma*; lam: *lambda*; nas: *nasion*; opi: *opisthocranion*; por: *porion*. The microchannels are concentrated along the main vault sutures and venous sinuses, except for a rhomboid-shaped distribution around the *bregma* (black arrows), associated with a low-density region in the anterior part of the parietal bones.

**Figure 4.**
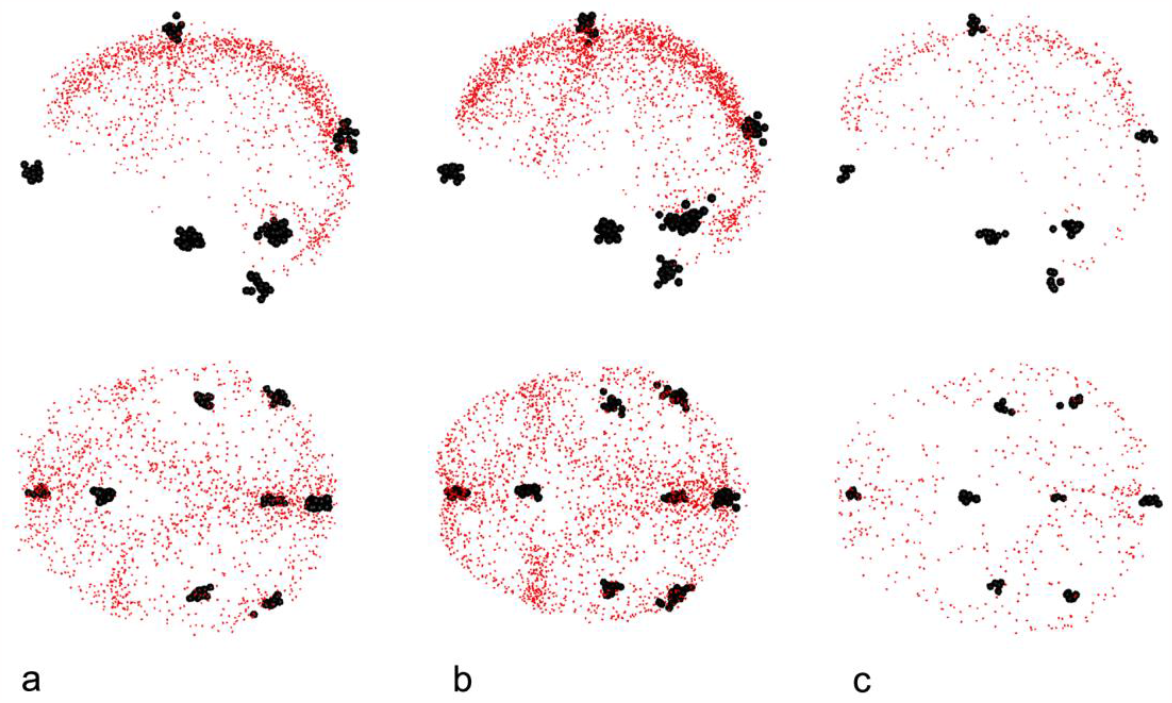
Maps of distribution on the microforamina in males (a), females (b), and juveniles (c) skulls, in left lateral and dorsal view.

## Discussion

Vascular functions are essential in brain biology, being in charge of oxygen supply, thermal exchanges, and hydrostatic support. The cerebral vessels redistribute energy, heat, and pressure within the brain volume, while the meningeal and diploic vessels manage the exchanges between the brain and skull. Craniovascular imprints are useful for investigating these vascular features in both living populations and extinct species, including osteological samples, radiographic data, and fossils (Píšová et al., 2017). In this context, microforamina can be particularly interesting because of their possible importance in thermal regulation (Zenker and Kubik, 1996). Bruner et al (2017) described these microchannels in a human parietal bone with more than 800,000 years, mostly around the lambdatic region. Hui and Balzeau (2023) mentioned these minor channels in Upper Paleolithic *Homo sapiens* and classic Neandertals, suggesting that both species display an overall similar distribution of these vessels. However, all these studies are limited to descriptive observations and, to date, these features are scarcely known even in modern human populations. In this article, we present the first quantitative assessment of these minor vascular channels, providing a comparative reference for further research and hypotheses on the variation and functions associated with these features.

### Number of microforamina

Most adult skulls display between 73 and 118 foramina (interquartile values). The skewed distribution (few smaller values and a tail of higher values fading until 190 spots) suggests that the lower threshold is probably associated with a functional limit. It remains to evaluate whether lower values could be associated with some pathological conditions, involving a loss of balance in oxygen supply, heat transfer, or drainage. All these structures are in fact involved in a fine balance between cerebrospinal fluid flow and thermal balance (Bertolizio et al., 2011; Adeeb et al., 2012, 2013). It would be also interesting to consider, in this sense, whether the relevant (functional) parameter is the overall number of channels or their local density.

Although adults display more microchannels than juvenile skulls, in the former group the number of microforamina seems not correlated to the age of the subject. This suggests that the increase of microchannels is associated with the early ontogenetic stages, reaching a stable value after maturity. This possibility is in agreement with the fact that also the diploic channels show an exponential increase just before adulthood (Rangel de Lázaro et al., 2020). It can be hypothesized hence that the two features (diploic channels and microforamina) share a proportional development, based on common factors. A match, in this sense, is likely, when considering that the microchannels probably connect the diploic network with the meningeal network.

Females display more microchannels (median = 106) than males (median = 88). Although the difference does not reach statistical significance, the p-value is however low. The sample size is not very large, so this lack of significance may be due to a lack of statistical power and small effect size (in this case, the effect size of this sample between females and males for the number of microforamina is 0.5). This sexual influence seems also supported by the fact that the only significant correlation between the number of microforamina and skull chords is associated with cranial length (*glabella*-*opisthocranion*), which is the most dimorphic diameters of the three considered here. The correlation is modest (R^2^=0.10) but, nonetheless, significant. Once corrected for cranial length, males and females display the same average number of channels, suggesting that the correlation is associated with the sexual difference. It remains to be considered whether the variation in the number of channels is primarily due to sex or cranial dimensions. The fact that shorter crania develop more channels seems counterintuitive, because the logical expectation is that an increase of surface should be associated with an increase of the vessels. Therefore, in this case, it is more likely to interpret this difference as a sexual trait, in which females develop more channels for reasons associated with sexual physiology and metabolism (including hormones, diploe development, etc.). This hypothesis is extremely interesting, and must be further investigated with additional samples.

### Distribution of microforamina

The map of distribution of the microchannels adds further information on their development. The channels are patently associated with the dorsal vault region, and their density decreases in the lower parts of the braincase. In particular, they are more concentrated along the sutures and the borders of the vault bone, than in the squamae. Such distribution may be due to specific local functions, or else be a secondary consequence of the diploe development, which is thicker in these areas. The relationship between diploic and vascular development is not straightforward because, although a threshold of thickness is necessary to allow the development of large vascular channels (Rangel de Lázaro et al., 2020), in adults the lumen size might not depend on the bone width (Eisová et al., 2016). Instead, the functional hypothesis gains interest when considering that the regions with high microvascular density spatially correspond with the course of major venous sinuses (in particular the superior sagittal and the sphenoparietal sinuses). This distribution strengthens further the functional hypotheses pointing to the reciprocal roles between the diploic channels and venous sinuses of the dura mater, in regulating both blood flow and the drainage of the cerebrospinal fluid. However, it is worth noting that, although the microchannels display a pronounced spatial correspondence with the venous sinuses, they apparently do not match the distribution of the main diploic channels. In fact, the larger diploic veins are generally described in the parietal squama, which instead presents a lower density of microforamina. The exception to the spatial correspondence between microchannels and venous sinuses is associated with the rhomboid pattern around the *bregma*, with a low density of vessels in the anteriormost region of the parietal bones. It can be speculated that this morphology might be associated with development issues. This region, actually, corresponds to the fontanellar space between the frontal and parietal bones that, occasionally, might generate an additional vault bone whose margins roughly resemble the distribution of the microforamina (Barberini et al., 2008). This morphology can be also coarsely recognized in our juvenile sample, and it can be considered whether it is associated with the different pace between bone and vascular development during early ontogeny.

### Limitations of the study

This first survey on the distribution and prevalence of endocranial microforamina has some general limitations that must be considered in the following studies. The first one concerns the use of standard tomography. Medical CT scans usually have a resolution of about 0.5 - 0.6 mm, which is sufficient for observing macroscopic craniovascular features (Lin & Alessio, 2009; Rangel de Lázaro et al., 2016, 2018). However, detecting foramina that are smaller than 0.5 mm is challenging, and can introduce a bias in the final count and localization. In young and juvenile specimens the detection of these channels is even more difficult, because of the scarce thickness of the diploe. In the case of osteological collections, as the one employed in this study, the state of conservation of some specimens can further hamper the recognition of these microchannels in the tomographic scans. Microtomographic analyses should be put forward to consider whether a finer resolution can improve the current results. Secondly, the current sample comes from a specific European population, and further research should consider a wider geographic variation. In this sense, it could be interesting to evaluate whether pronounced cranial differences, climate, or pathological conditions, can be associated with variations of these vascular features. In this case, although the results presented here look consistent, larger samples can also improve the statistical power, confirming and evidencing finer differences between sexes. Lastly, the ontogenetic changes have been here only introduced with a preliminary sample. Dedicated research on this topic is indeed mandatory.

## Conclusions

To our knowledge, this is the first study describing and quantifying the frequency and distribution of endocranial microforamina in humans. Results support information in terms of both quantity (the amount of channels) and quality (the distribution of the channels). On the one hand, this information is necessary to consider whether vascular morphology underwent evolutionary changes associated with the process of encephalization in hominids. At the same time, this same information can be relevant in medicine and health sciences, when considering the possible importance of these features in brain thermal regulation (Zenker and Kubik, 1996). More recently, it has been proposed that these microvessels are involved in the migration of immune cells from the vault bones to the brain, after cerebral damage and inflammation (Herisson et al., 2018). Myeloid cells would be produced as hematopoietic response in the bone marrow, then flowing into the cerebral tissues through these microchannels, to repair and manage the damaged or inflamed tissues. Such a generalized mechanism is activated in many distinct pathological conditions, ranging from stroke to Alzheimer’s disease. We hope that this first anatomical survey on these traits will call attention to the contribution of craniovascular morphology and morphogenesis on brain biology, an issue that is probably still neglected in evolutionary anthropology and biomedicine.

## Acknowledgments

EB is funded by Project PID2021-122355NB-C33 (MCIN/AEI/10.13039/501100011033/ FEDER, UE), and by the Italian Institute of Anthropology (ISITA). SE is funded by the Ministry of Culture of the Czech Republic (grant number DKRVO 2024-2028/7.I.a, 00023272, National Museum). We would like to thank the Department of Anthropology and Human Genetics of Charles University in Prague for providing the CT sample of the Pachner collection. The authors declare no conflict of interest.

## Data Availability Statement

Data are available under request.

## Notes

### Competing Interest Statement

The authors have declared no competing interest.

